# PhageTerm: a Fast and User-friendly Software to Determine Bacteriophage Termini and Packaging Mode using randomly fragmented NGS data

**DOI:** 10.1101/108100

**Authors:** Julian Garneau, Florence Depardieu, Louis-Charles Fortier, David Bikard, Marc Monot

**Affiliations:** Institut Pasteur, Laboratoire Pathogenèse des bactéries anaérobies, Département de Microbiologie, Paris, France.; Département de Microbiologie et d'infectiologie, Faculté de Médecine et des Sciences de la Santé, Université de Sherbrooke, Sherbrooke, QC, Canada.; Institut Pasteur, Groupe Biologie de Synthèse, Département de Microbiologie, Paris, France.; Université Paris Diderot, Sorbonne Paris Cité, Paris, France.

**Keywords:** Next-generation sequencing, NGS, Bacteriophage, Termini, Packaging

## Abstract

Bacteriophages are the most abundant viruses on earth and display an impressive genetic as well as morphologic diversity. Among those, the most common order of phages is the Caudovirales, whose viral particles packages linear double stranded DNA (dsDNA). In this study we investigated how the information gathered by high throughput sequencing technologies can be used to determine the DNA termini and packaging mechanisms of dsDNA phages. The wet-lab procedures traditionally used for this purpose rely on the identification and cloning of restriction fragment which can be delicate and cumbersome. Here, we developed a theoretical and statistical framework to analyze DNA termini and phage packaging mechanisms using next-generation sequencing data. Our methods, implemented in the PhageTerm software, work with sequencing reads in fastq format and the corresponding assembled phage genome.

PhageTerm was validated on a set of phages with well-established packaging mechanisms representative of the termini diversity: 5’cos (lambda), 3’cos (HK97), pac (P1), headful without a pac site (T4), DTR (T7) and host fragment (Mu). In addition, we determined the termini of 9 *Clostridium difficile* phages and 6 phages whose sequences where retrieved from the sequence read archive (SRA).

A direct graphical interface is available as a Galaxy wrapper version at https://galaxy.pasteur.fr and a standalone version is accessible at https://sourceforge.net/projects/phageterm/.

## INTRODUCTION

Phages, the viruses of Bacteria, come in a diversity of shapes and sizes^1^. They all produce viron particles consisting of a protein shell, with in some cases the addition of a membrane. Their nucleic acid content can be double stranded DNA (dsDNA), single stranded DNA, double stranded RNA or single stranded RNA. Different conformations of the nucleic acid also exist with some phage genomes being encapsidated as circular and other as linear molecules. The vast majority of phages described to date produce capsids containing linear dsDNA, but even when considering these phages only, an impressive diversity of packaging mechanisms has been described, leading to various DNA termini^2^.

Towards the end of their infection cycle, dsDNA phages generally form concatemers which are cut by the terminase during packaging to form the mature chromosome^3^. One can distinguish four main mechanisms used by phages to initiate and then terminate packaging while recognizing their own DNA rather than the host DNA: (i) The terminase can recognize a specific site where it introduces a staggered cut (cos site), generating fixed DNA termini with cohesive ends that can either have 5’ or 3’ overhangs (e.g. lambda^4^, HK97^5^). (ii) A fixed position can be recognized on the phage DNA where direct terminal repeats (DTR) will be generated by extension synthesis at the 3’ ends of staggered nicks. The size of these DTRs can range from just over a hundred bases (e.g. T3^6^, T7^7^) to several thousand bases (e.g. T5^8^). Phage N4 carries terminal repeats with an accurate terminus on the left end but several possible termini on the right end^9^. (iii) The terminase can initiate packaging on the phage concatemer at a specific packaging site (pac site), but following cuts are made when the phage head is full at variable positions (e.g. P1^10^, P22^11^). This leads to capsids containing circularly permutated genomes with redundant ends used to circularize the phage genome through recombination after injection in the host cell. (iv) T4-like phages use a variant of this headful packaging mechanism in which no pac site is recognized, but packaging is rather initiated randomly^12^. These phages will usually degrade the host DNA, ensuring that only their own DNA is packaged.

One should also add to these, three less frequent packaging strategies that do not involve the formation of concatemers: (i) Phage P2 carries a cos site but the packaging substrate is circular dsDNA^13^ (ii) Phage Mu replicates through transposition in the host genome and carries pieces of the host DNA as its termini^14^. (iii) The Bacillus phage phi29 carries covalently bound proteins at its DNA termini^15^.

When considering the small number of phages for which the termini was precisely studied, it is likely that yet other packaging mechanisms exist in nature. In this study we investigate how the information gathered by high throughput sequencing technologies, and in particular Illumina technologies, can be used to determine the DNA termini and packaging mechanisms of dsDNA phages. The wet-lab procedures traditionally used for this purpose rely on the identification and cloning of restriction fragment which can be delicate and cumbersome, in particular in the case of circularly permutated phages where the packaging sites will be present in sub-stoichiometric bands after digestion^16^. Additional work might also have to be performed to retrieve the packaging orientation, the exact position of the termini and the cohesive sequence.

Many high throughput sequencing methods rely on the random fragmentation of DNA followed by DNA ends repair and adapters ligation. After the fragmentation process, natural DNA termini will be present once per phage genome, while DNA ends produced by fragmentation will fall at random positions along the genome. This means that many more reads will start at the phage termini than elsewhere in the genome. This observation was made by several groups, and was recently used by Li *et al.*^17^ to characterize the DNA termini of several phages.

We propose here a theoretical and statistical framework to analyze DNA termini and phage packaging mechanisms using next-generation sequencing data. The software was validated on a set of phages with well-established packaging mechanisms representative of the termini diversity: 5’cos (lambda), 3’cos (HK97), pac (P1), headful without a pac site (T4), DTR (T7) and host fragment (Mu). In addition, we applied our methods to decipher the termini of 9 *Clostridium difficile* phages and 6 phages whose sequences where retrieved from the sequence read archive (SRA). Our methods are implemented in the PhageTerm software which we are making freely available. For direct graphical interface operation, a Galaxy wrapper version is also available at https://galaxy.pasteur.fr.

## METHODS

### Phage DNA extraction and sequencing

#### Escherichia coli phages

Phage lambda, HK97, T7, P1 and T4 were propagated on E. coli strain MG1655 in LB supplemented with CaCl2 (5 mM). A thermosensitive variant of bacteriophage Mu (MuCts62) was induced from strain RH8816, a derivative of strain MC4100^18^. Strain RH8816 was grown at 30°C to OD~0.5 and induced at 42°C during 2 hours.

Phage capsids were isolated using a PEG precipitation step followed by cesium chloride (CsCl) gradient purification. First, solid NaCl was added to the phage lysate to a final concentration of 1M and incubated on ice for 1h, followed by centrifugation at 11,000g for 10 min at 4°C to remove the cellular debris. PEG8000 was added to a final concentration of 10% w/v for 1h on ice followed by centrifugation at 11,000g for 10 min at 4°C. The pellet was resuspended in PBS (1X) and incubated for 1h at room temperature. An equal volume of chloroform was then added followed by centrifugation at 3000g for 15 min at 4°C. The aqueous phase which contains the phage particles was recovered and further purified by ultracentrifugation with a gradient of cesium chloride (CsCl). A volume of 3 ml of each of two cesium chloride solutions (density of 1.3 g/ml- 0,41 g/ml of water and density of 1.5 g/ml- 0,68 g/ml of water) and a volume of 2ml of a CsCl solution (density of 1.6 g/ml- 0,82 g/ml of water) was added sequentially from more to less dense with a pipette into the bottom of a polyallomer tube. Phage suspensions up to 3 ml were then layered on the top of the gradient, and tubes were centrifuged at 100,000 g for two hours in a Beckman SW41 swinging bucket rotor. During this time, a roughly linear gradient of CsCl was formed in the tubes. Phages concentrated as a visible band were collected after puncturing the tube with a needle under the corresponding band. A second step of equilibrium ultracentrifugation in cesium chloride solution (density of 1.5 g/ml) was performed at 150,000 g with Beckman SW60 rotor. Phages were then dialyzed twice against water for 20 min and overnight against Tris 10 mM and NaCl 150 mM. Concentrated samples of phage lysates were treated with DNase and RNase for 30 min at 37°C, DNase was inactivated by EDTA (pH 8.0, 5 mM) for 10 min at 65°C and then proteinase K (0,5 mg/ml) and SDS (0,5 %) was added for 30 min at 55°C. Phage DNA was then purified using a typical protocol with phenol/chloroform/isoamylic alcohol followed by precipitation with acetate sodium (0,3M) and ethanol 100%. Pellets were washed with ethanol 70% and resuspended in TE (1X). Phage DNA was sequenced using the TruSeq library preparation kit on a HiSeq device.

#### Clostridium difficile phages

The NGS sequences of 8 Clostridium phages are reported: three temperate *Siphoviridae* phages (phiCD24-1, phiCD111, and phiCD146), and five temperate *Myoviridae* phages (phiCD481-1, phiCD505, phiCD506, phiMMP01 and phiMMP03). Phage DNA was isolated from 10 mL of purified phage lysate using the QIAGEN Lambda Mini Kit as described previously^19^. Since no suitable propagating host could be identified, we extracted phage DNA directly from a crude induction lysate. The quality of the DNA preparations was verified on agarose gel and the absence of contaminating bacterial DNA was verified by PCR using primers targeting the triose phosphate isomerase gene^20^. Single-end multiplex TruSeq libraries were created according to Illumina’s specifications. Sequencing was performed on the Illumina HiSeq 2000 platform of the Pasteur Institute (Paris, France). The 110 bp sequencing reads (12 to 19 Mb, >1000X) were assembled into a single contig for each phage using Velvet^21^. Finally we used the MicroScope workflow^22^ to generate an automatic functional annotation for putative coding sequence (CDS), which was then manually curated.

#### Phage *de novo* assembly

To find the termini of an unknown phage, the first step consisted of a *de novo* assembly using NGS raw data. To validate that the assembly did not alter the result of the software, we assembled the 5 reference phages using spades^23^ with standard options. Assemblers will frequently introduce mistakes at the edges of contigs due to the presence of low abundance reads that do not correspond to the actual phage sequence. This can usually be seen as a drop in coverage at the contig ends. Phage assembly genome are available in the Sourceforge deposit (https://sourceforge.net/projects/phageterm).

#### Nucleotide sequence accession numbers

The complete genome sequences of phiCD24-1, phiCD111, phiCD146, phiCD481-1, phiCD505, phiCD506, phiMMP01 and phiMMP03 were deposited in European Nucleotide Archive under the accession no. LN681534, LN681535, LN681536, LN681538, LN681539, LN681540, LN681541, LN681542. The *de novo* assemblies performed on reference phages are available at Sourceforge (https://sourceforge.net/projects/phageterm). Phages used in this study are described in Table S1. The raw reads data of all phages was deposited in sequence reads archives (SRA) under the accession SRP093616 (Table S2).

### Software implementation

PhageTerm is a standalone program designed for biologists which purpose is to perform termini and packaging mode analysis for any bacteriophage of interest. PhageTerm is written in Python 2.7, the source code and documentation are freely available at https://sourceforge.net/projects/phageterm under a GPL license (GPL 3.0). PhageTerm is compatible on three operating systems (Linux, Mac OS X, and Windows) in command line (require Python 2.7.X with the following library: matplotlib 1.3.1, numpy 1.9.2, pandas 0.19.1, sklearn 0.18.1, scipy 0.18.1, statsmodels 0.6.1 and reportlab 3.3.0.). For direct graphical interface operation, a Galaxy wrapper version is also available at https://galaxy.pasteur.fr.

### Inputs and process

PhageTerm works with two mandatory inputs: (i) a file containing the phage sequence in Fasta format (Reference Sequence); and (ii) a file containing the sequencing reads in fastq format (Reads File). Optional inputs can also be chosen: the phage name to be used as an output prefix (default: Phagename), the length of the seed used for reads in the mapping process (default: 20), a surrounding value for peaks merging (default: 20), a file containing paired-end reads in fastq format (Paired File), the host genome sequence in fasta format (Host Sequence) and the number of cores you want to use.

### PhageTerm Analysis

Reads are mapped on the reference to determine the starting position coverage (SPC) as well as the coverage (COV) in each orientation. When paired-end information is provided, the fragment coverage is computed by considering that all the bases between a pair of reads are covered, otherwise if single reads are provided, sequence coverage is computed as an approximation of fragment coverage. These values are then used to compute the variable X = SPC / COV. This variable has several interesting properties that are useful to determine DNA termini.

#### Cos phages

The coverage at position i is determined by the number of fragments that start at position i plus the number of fragments that start before i and cover i. In the case of a fixed DNA terminus no reads should start before the terminus and therefore X=1. Note that different outcomes are expected for cos phages with 5’ or 3’ cohesive ends. The end-repair enzymes typically used during sequencing library preparation will fill 5’ overhangs and chew 3’ overhangs. A value of X=1 is thus indeed expected for 3’ cos phages. However, end-repair will produce overlapping ends for 5’ cos phages: fragments that end at the right terminus will be seen as covering the left terminus and conversely. The expected value of X is thus 0.5. Note however that when single reads are used, sequence coverage will be computed instead of fragment coverage, which can affect the expected value of X. Indeed, if the average fragment length is smaller than the read length, then little to no forward reads will be obtained that start before the left terminus and cover it. As a consequence, the expected value of X at the left terminus will be 1. The same argument holds in the other orientation.

#### DTR phages

In the case of DTR phages, for N phage particles in a sample that undergo fragmentation, there should be N fragments that start at the terminus, and N fragments that cover the edge of the repeat on the other side of the genome. As a results X is expected to be 0.5. Similarly to what is described above for 5’ cos phages, the expected value of X can change if single reads are used and the sequence coverage is computed instead of fragment coverage. This is especially true for short DTR. Indeed when the read length is smaller than the average fragment length minus the DTR size, then little to no forward reads will be obtained that start before and cover the edge of right repeat. The same argument also holds in the other orientation. As a result a value of X as high as 1 can be obtained.

#### Pac phages

In the case of pac phages, for N phages present in the sample, there should be N/C fragments starting at the pac site, where C is the number of phage genome copies per concatemer. In the same sample N fragments should cover the pac site position. Therefore, at the pac site position X is expected to be (N/C)/(N+N/C) = 1/(1+C).

#### Calling significant termini

Another interesting property of X is that its average value at positions along the genome that are not termini is expected to be 1/F, where F is the average fragment size which can be easily computed from paired-end sequencing data. Indeed, if the average number of reads that start at a random position is R, then the average number of fragments that cover the same random position is R*F. Note that 1/F should always be much smaller than the expected value of X at termini position in any of the situations described above. Indeed the average fragment size in illumina sequencing libraries usually ranges from 300-800bp. This should also be true for pac sites. Indeed, 1/(1+C) >> 1/F since the number of phage genome copies per concatemer (C) reported in the literature is typically smaller than 10^24^. Nonetheless, calling the position of pac sites can be made more difficult because of the experimental noise introduced in the many steps of library preparation. To assess whether the number of reads starting at a given position along the genome can be considered a significant outlier, PhageTerm first segments the genome according to coverage using a regression tree. A gamma distribution is then fitted to SPC for each segment and an adjusted p-value is computed for each position. A significance threshold of 0.01 was chosen to consider that a peak in X represents a valid terminus. Finally if several peaks with an adjusted p-value lower than 0.05 are detected within a small sequence window (default: 20bp), the position is deemed significant. This allows detecting packaging sites for which the terminase can cleave at several nearby positions. In such cases the number of reads starting at nearby peaks is added for subsequent analysis, and the position of the highest peak is reported.

#### Phage classification

The position of termini can be fixed (X>0.6), multiple (several significant peaks, X>0.6), preferred (a single significant peak, X>0.6) or multiple-preferred (several significant peaks, X>0.1). Note that “multiple” can only occur if the phage genome is broken into several chromosomes. To our knowledge there are no report of such dsDNA phages. In the case of a preferred peak on one strand, we look for an increased coverage (>10% of the average coverage) after the peak within 2% of the genome length. Such an increased coverage is expected for *pac* phages. In this case the termini on the other strand is said to be distributed. When no significant peak is detected the terminus is classified as random.

The following rules are then applied to classify the phages: When significant peaks with X>0.1 are found on both strands, the distance between the peaks is used to differentiate cos phages and DTR phages. The largest cohesive ends described to our knowledge are 19 bases^16,25^, and the smallest DTR, 131 bases from phage T3. We therefore placed a threshold at 20bp. An additional criterion to classify a phage as DTR is that the coverage between the peaks should be at least 10% higher than the average coverage. Phages classified as cos are determined to have 5’ overhangs if the forward terminus is to the left of the reverse terminus, and 3’ overhangs otherwise. When a significant peak with X>0.1 is found only on one strand, the phage is considered to be a pac phage. PhageTerm also provides the orientation of packaging for pac phages, as determined by the strand carrying the significant peak.

Phages for which no significant peaks are found could either use a headful packaging mechanism without a preferred packaging site (T4-like), or be Mu-like phages. To differentiate both possibilities, the software computes the number of fragment for which one read matches the host and the other read matches the phage genome. Phages in the sample will on average be fragmented into G/F pieces where F is the average fragment size and G the phage genome size. In the case of Mu-like phages, out of G/F phage fragments, two should be hybrid fragments. The proportion of hybrid fragments is thus expected to be 2*F/G. Note that these fragments will only be detected if the extent spanning the host genome and the extent spanning the phage genome are both longer than the seed sequence (S) used to align the reads. The theoretical expectation of the proportion of hybrid fragments can thus be corrected as 2*(F-2*S)/G. PhageTerm assigns the Mu-like class if the proportion of hybrid fragments is at least half the expected value. The position of hybrid reads on the phage genome is then used to estimate the termini positions (Figure 3). Finally phages for which no hybrid fragments are found are classified as unknown.

Lower than expected X values could indicate that the phage DNA is contaminated with unpackaged DNA, or that problems occurred during the sequencing library preparation. If these problems can be excluded, X values that deviate from the expectations might point towards novel packaging mechanisms.

#### Implementation of the method from Li *et al.*

The second approach is based on the calculation and interpretation of three specific ratios R1, R2 and R3 as suggested in a previous publication from *Li et al.*^17^. The first ratio, is calculated as follow: the highest starting frequency found on either the forward or reverse strands is divided by the average starting frequency, R1 = (highest frequency/average frequency). Li’s *et al.* have proposed three possible interpretation of the R1 ratio. First, if R1 < 30, the phage genome does not have any termini and is either circular or completely permuted and terminally redundant. The second interpretation for R1 is when 30 ≤ R1 ≤ 100, suggesting the presence of preferred termini with terminal redundancy and apparition of partially circular permutations. At last, if R1 > 100 that is an indication that at least one fixed termini is present with terminase recognizing a specific site. The two other ratios are R2 and R3 and the calculation is done in a similar manner. R2 is calculated using the two highest frequencies (T1-F and T2-F) found on the forward strand and R3 is calculated using the two highest frequencies (T1-R and T2-R) found on the reverse strand. To calculate these two ratios, we divide the highest frequency T1 by the second highest frequency T2. So R2 = (T1-F / T2-F) and R3 = (T1-R / T2-R). These two ratios are used to analyze termini characteristics on each strand taken individually. *Li et al.* suggested two possible interpretations for R2 and R3 ratios combined to R1. When R1 < 30 and R2 < 3, we either have no obvious termini on the forward strand, or we have multiple preferred termini on the forward strand (when 30 ≤ R1 ≤ 100). If R2 > 3, it is suggested that there is an obvious unique termini on the forward strand. The same reasoning is applicable for the result of R3, used to analyze the termini on the reverse strand. Combining the results for ratios found with this approach, it is possible to make the first prediction for the viral packaging mode of the analyzed phage. A unique obvious termini present at both ends (both R2 and R3 > 3) reveals the presence of a cos mode of packaging. The headful mode of packaging PAC is concluded when we have a single obvious termini only on one strand.

Some additional information could also be extracted depending on phage characteristics i) an estimation of phage concatemers; and ii) the sequence located between the two termini, for the specific case where the phage have defined termini (fixed or preferred) or recognized as a cos (sequence length under 20 bases are considered as cohesive ends, others as direct repeats). The output of the two methods are consolidated in a detailed PDF report produced at the end of the analysis pipeline. Other outputs are also provided: statistics table of each position (csv format), cohesive ends or direct terminal repeats (fasta format) and the phage genome sequence reorganized following termini and completed with extremity or repeats if needed.

### Performance

The PhageTerm software was optimized to use multiple core and to minimize the memory consumption. During the whole process, the memory footprint remained low (< 1Gb). In our experiments, PhageTerm processed reads at a rate of more than 1 million reads per CPU-hour (3.5GHz Intel Xeon E5 with 12MB L3 cache).

### How inputs affect results ?

#### Reference phage genome

The reference phage genome could contain its direct terminal repeats depending on the method used for its assembly. In this case, PhageTerm should detect two peaks for each strand. We test this hypothesis using the T7 reference genome (NC_001604, Table S1) and the PhageTerm output was multiple peaks on both strand for the two analysis methods (T7-reference report in the sourceforge deposit, https://sourceforge.net/projects/phageterm).

#### NGS reads quantity

To test the reliability of the two methods, results were analysed with a decrease number of reads (Table S3). The majority of the PhageTerm results are conserved up to 10K reads and only *C. difficile* phages needed up to 100k reads. The software analysis methods are very resilient to low quantity sequencing.

## RESULTS

Different mechanisms of DNA packaging can lead to a variety of DNA ends (Table S4). When DNA ends occur at fixed or preferred positions along the phage genome, we can expect to observe more reads starting at these positions than elsewhere in the genome (Figure 1A). We sequenced 5 well studied *E. coli* phages encompassing various packaging mechanisms (Lambda, HK97, P1, T4, Mu) in order to validate our strategy. The PhageTerm analysis aligns the reads to a reference sequence provided by the user and computes the number of reads starting at each position (SPC), the sequence coverage (COV) and a signal value X described in the materials and methods and which simply corresponds to SPC/COV. This X variable displays several useful properties that allows classifying the phages.

**Figure 1.**
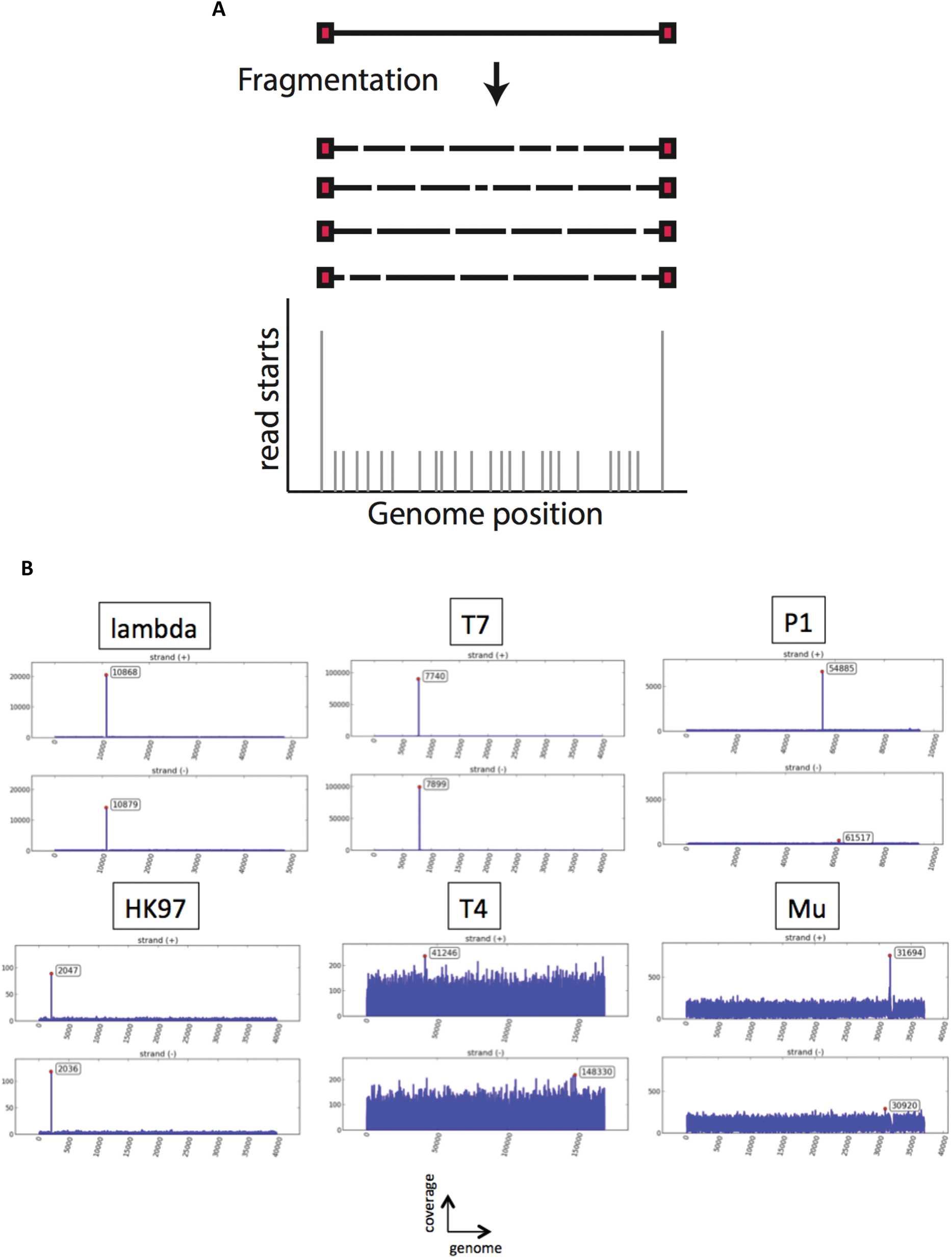
Biases in the number of reads starting at natural DNA ends. A) Random fragmentation of DNA followed by adapted ligation and sequencing leads to biases in the number of reads that start at the phage termini. The black line represents the phage genome, the red squares are the phage termini. After the fragmentation process, natural DNA termini will be present once per phage genome, while DNA ends produced by fragmentation will fall at random positions along the genome. B) Starting position coverage for each strand of 6 reference phages (lambda, HK97, T7, P1, T4, Mu).

## Fixed DNA ends

### COS

Cos phages are expected to display a single peak in each orientation with a values of 0.5<X<1 (see materials and methods). These phages thus provide a very strong signal and can be easily identified. If the phage DNA sample was not contaminated with unpackaged DNA, then no reads should be recovered that cross the cos site. As a result, *de novo* assemblers should naturally place the DNA ends at the contig limits. Note that different results are expected for cos phages with a 5’ overhang and with a 3’ overhang (Figure S1). The enzyme typically used during the step of DNA end repair will fill 5’ overhangs but chew 3’ overhangs. Phages with 5’ overhangs are expected to share the same terminal sequence over the length of the cohesive end, while phages with 3’ overhangs will have different terminal sequences. In practice, reads are frequently recovered that do cross the cos site, which can lead the assembler to place the cos site at a random position along the contig. In this case the peak positions will fall in close proximity to each other. On the one hand, cos phage with 5’ overhangs will have the forward peak positioned to the left of the reverse peak with the region in between showing twice the average sequence coverage (Figure 2A). On the other hand, cos phages with 3’ overhangs are expected to have the forward peak to the right of the reverse peak, and the sequence in between is expected to have a very small coverage (Figure 2B).

**Figure 2.**
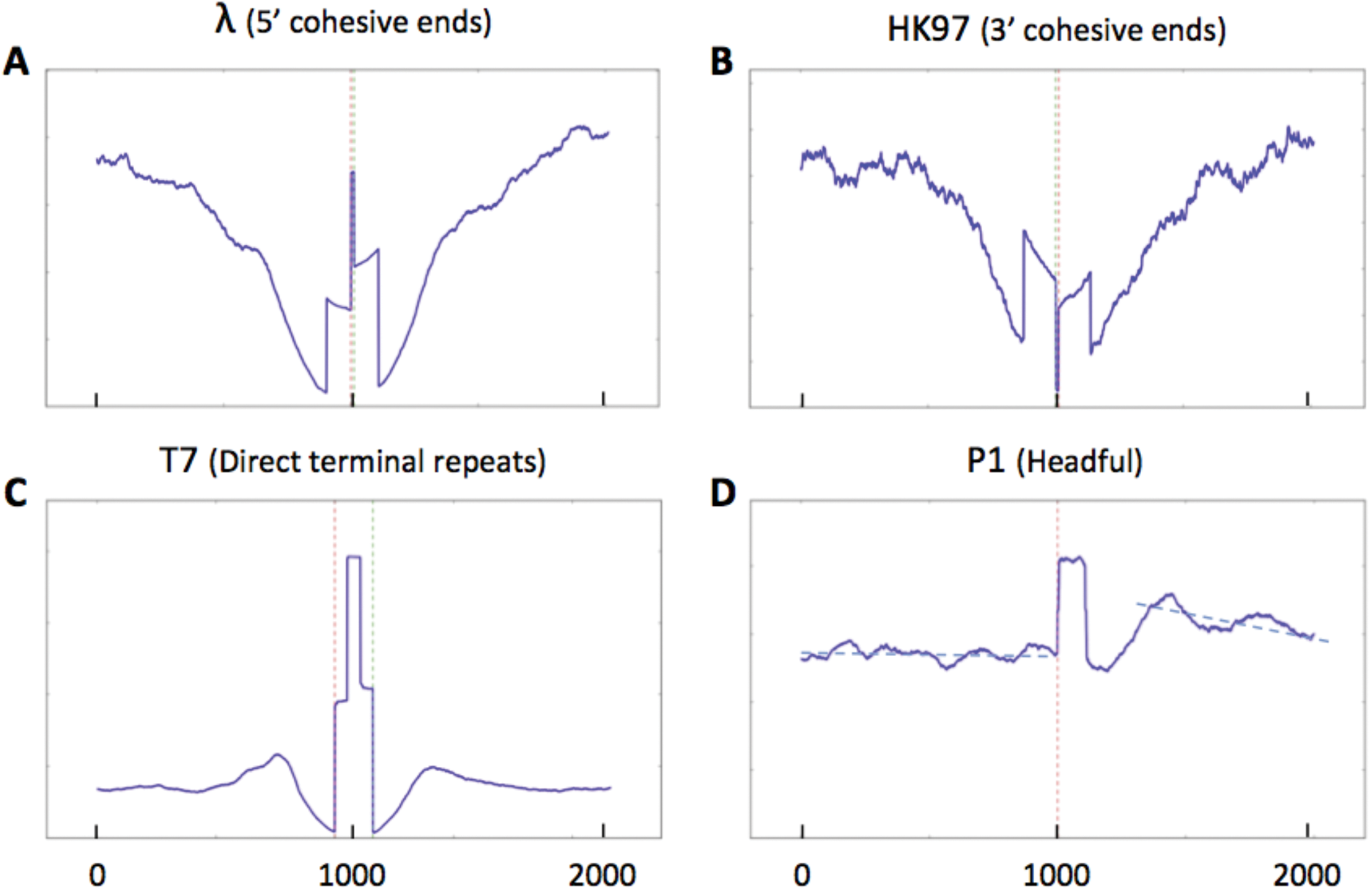
Sequence coverage at termini position. The sequence coverage around the termini identified by PhageTerm was plotted for the following phages: lambda, HK97, T7 and P1. Exact termini positions are represented by dotted red line (Red: left; Green: right). Note the higher coverage obtained for phage P1 after the packaging site.

As an example of a 5’ cos phage we sequenced the well-studied siphoviridae coliphage lambda^26^. The virion genome is known to have two 5’ cohesive single-stranded ends of 12 bases. For 3’ cos phage, HK97^5^ was sequenced. The PhageTerm software was able to determine the exact characteristics of lambda and HK97: the nature of the termini (Figure 1B), the cos mode of packaging and the length of their cohesive sequence (Table 1). Phage Efm1 from Enterococci was described as a 3’ cos phage^27^, and was also accurately recognized by PhageTerm. Among Clostridium phages that we sequenced, PhiCD481-1 and PhiCD506 also matched the theoretical description of cos phages with 3’ overhangs (Figure 2B).

**Table 1.** Characteristics of reference phages.

| Phage Name | Genome (G) | Left (X) | Right (X) | Class | T/R | Fragment size (F) |
| --- | --- | --- | --- | --- | --- | --- |
| Lambda | 48 kb | 0.59 | 0.41 | COS (5') | 126 | 421 |
| HK97 | 39 kb | 0.51 | 0.60 | COS (3') | 115 | 353 |
| T7 | 39 kb | 0.47 | 0.51 | DTR (short) | 594 | 468 |
| P1 | 94 kb | 0.20 | - | Headful (pac) | 101 | 494 |
| T4 | 168 kb | - | - | - | - | 463 |
| Mu | 36 kb | - | - | Mu-like | - | 485 |

#### DTR

Phages with direct terminal repeats are expected to have one nice peak in each orientation, and the forward peak should be to the left of the reverse peak. The region in between the peaks should show twice the average sequence coverage (Figure 2C). Because of their terminal redundancy, DTR phages will typically be attributed a random first base by DNA assemblers. A typical example of this is phage T7, a podoviridae that has exact short direct terminal repeats of 160 bases^25^. The PhageTerm software was able to determine the exact characteristics of T7: nature of its termini (Figure 1B), the termini positions and the exact length of its direct terminal repeats (Table 1).

### Headful packaging

Phages using a headful packaging mechanism will typically first produce a concatemer containing several copies of their genome. During packaging a first cut is made at the packaging site (pac site), but following cuts are made when the phage head is full at variable positions. The expected value of X can be computed to be 1/(C+1), where C is the number of phage genome copies per concatemer (see material and methods). Because packaging is directional and no precise cut is made upon termination of packaging, a peak is expected only in a single orientation, which also informs us about the direction of packaging. In the region after this peak, where the second cut is made, a slight increase in coverage is expected, as part of this region will be present twice in many phage particles. Note that the signal obtained to determine the position of the pac site is C+1 time weaker than that for cos phages (Figure S3). This phenomenon is amplified by the fact that the cleavage position of the terminase at pac sites can be imprecise, leading to several possible termini. As an example of this packaging mechanism we sequenced phage P1, a myoviridae enterobacteria phage. The PhageTerm software was able to determine the exact characteristics of P1: nature of the termini (Figure 2D), the *pac* mode of packaging and an estimation of the number of concatemers.

Other phages have been described to package their genome through a headful packaging mechanisms but with no preferred packaging signal. The packaged phage genomes are circularly permuted with random termini. No signal can be recovered for this type of phage. As an example we sequenced T4, a myoviridae enterobacteria phage (Figure 1B). These phages will typically destroy the host cell DNA before packaging ensuring that only the phage DNA is packaged.

### Mu-like phages

Mu-like phages are temperate phages that amplify their genome through replicative transposition. During packaging, a first cut is made at a given distance from the phage end in the host genome. Packaging then proceeds through a headful mechanism with the second cut being made on the other side of the phage in the host DNA. The packaged DNA termini thus corresponds to various fragments of the host DNA (Figure S4). A specific analysis is performed to discover this type of packaging mechanism which relies on paired-end sequence information. We look for hybrid fragments, i.e. pairs of reads where one read matches the phage genome and the other read the host genome. The presence of a substantial amount of such fragments strongly points towards this mode of replication and packaging.

We sequenced phage Mu as the father of this category. The PhageTerm software results are in line with those expected: no significant peak in the value of X were found (Figure 1B), but a high number of hybrid inserts were detected (Figure 3). The host termini of Mu are asymmetric, a short fragment at one side (~50) and a long one at the other side (~2000bp). Unfortunately, the short read length provided by illumina sequencing does not allow to determining these fragment sizes easily and systematically.

**Figure 3.**
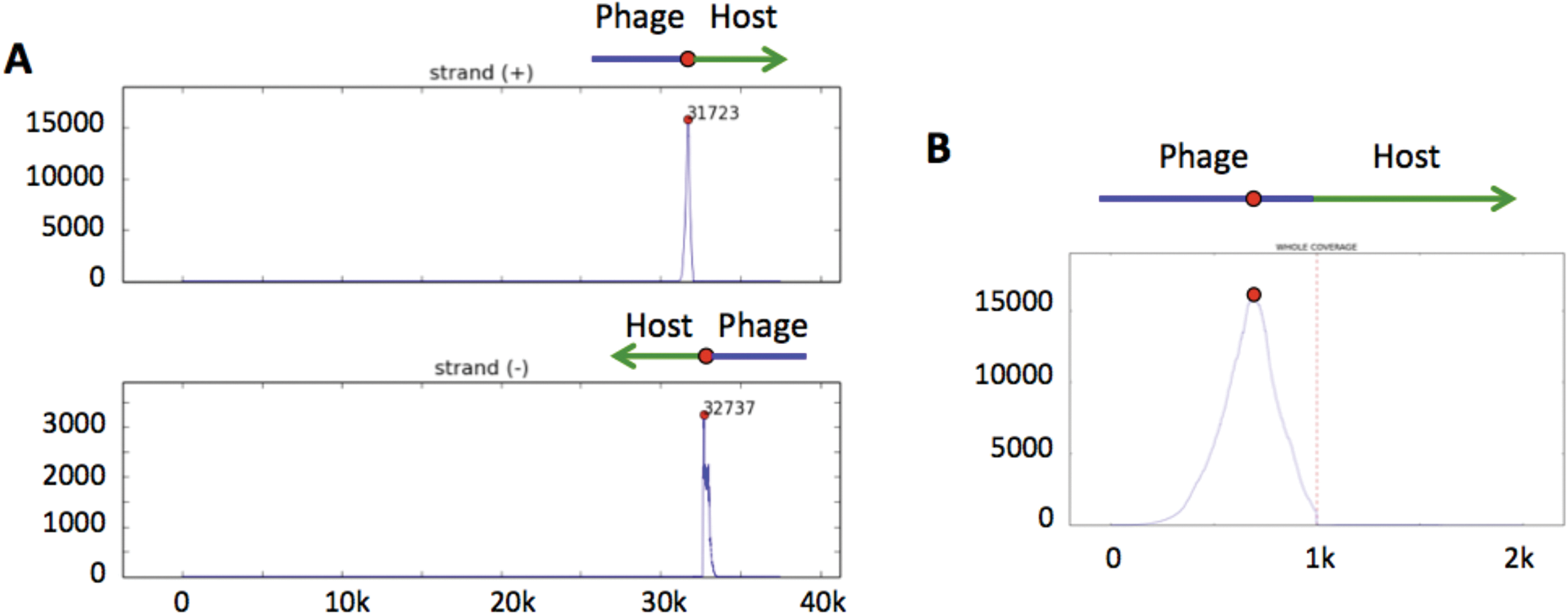
Detection of Mu-like phages. Hybrid fragments with one read on the phage genome and one read on the host genome are detected. A) The sequence coverage of reads belonging to hybrid fragments of phage Mu is plotted along the phage genome. B) Zoom in around the right terminus of phage Mu. The dotted red line represents the terminus position.

### Peak size

Another interesting variable to consider when analysing phage sequencing data is number of reads starting at the termini (T) divided by the average number of reads starting at random positions (R). For phages with fixed termini, the size of the signal defined as T/R (R1 in Li *et al.*^17^) is expected to be equal to the average fragment size (F). Deviations from this expectation might indicate inefficient ligation of the sequencing adapters to the phage DNA termini, or the presence of other possible DNA termini. Sequencing results for phage T7 are in good agreement with this predictions. However, T/R is markedly smaller than expected for phages lambda and HK97 (Table 1). This is presumably due to an inefficient repair of the cohesive ends during our sequencing library preparation.

In the case of *pac* phages, an interesting observation is that knowing T/R and F, it is possible to estimate the number of phage copies per concatemer: C = 1/X - 1 = F*R/T - 1. For phage P1 we can estimate the number of phage copies per concatemer to be ~4, in good agreement with previous estimates^10^.

### PhageTerm results

The PhageTerm software produces a detailed report containing several useful plots (coverage, SPC), a data table, information about termini types and packaging mode. As an output, a re-centered sequence file is generated. Some additional information is also extracted depending on phage characteristics: i) an estimation of the phage concatemers size (pac phages); ii) the sequence located between the two termini (cos and DTR).

#### Clostridium difficile Phages

*Clostridium difficile* is currently the principal cause of antibiotic-induced infectious diarrhea in humans^28^. Most *C. difficile* isolates analyzed to date carry one or several prophages, however only a few of them have been fully characterized. The packaging mode of four phages included in this study have been determined experimentally in our laboratory using the ligation-digestion method and could be used to compare the results we obtain with the software. PhageTerm determined PhiCD481-1 and PhiCD506 to be cos phages, and found 13nt 3’ overhangs identical for both phages (Table S1). These phages also showed the expected decrease in coverage between the termini of 3’cos phages (Figure 2B). PhiCD146, PhiMMP01 and PhiMMP03 were found to be pac phages. PhiCD211 was found to have direct terminal repeat of 378 bp. No termini could be detected for PhiCD505 and PhiCD24-1, which could indicate a T4-like packaging mechanism. Finally phiCD111 showed a weird coverage plot, with decreasing coverage after a significant peak at the end of the genome, and another significant peak at another distant position. No packaging mechanism could be assigned to this phage.

#### SRA Phages

To test the versatility and robustness of the PhageTerm software on data generated by others, we have fetched the raw sequencing data of six phages (T3, T7, HP1, Efm1, Slur09 and PBES-2) from the sequence read archive (SRA). Among these phages, four have known termini and packaging mode (T7, T3, HP1 and Efm1) detailed in table S1. The PhageTerm software was able to identify the right termini and packaging mode for T3 and T7 along with an exact short DTR of respectively 160 and 231 bases. For phage Efm1 and PBES-2, the PhageTerm software found cohesive sequence of 9 bases and direct repeats of 443 bases, respectively.

#### Important note on sequencing library preparation methods

The methods described here rely on the random fragmentation of the phage DNA and the availability of DNA termini to adapter ligation. As a consequence methods that rely on transposases to ligate the adapters, such as Nextera, should not produce suitable data for PhageTerm and similar strategies^29^. We confirmed this by analyzing the data of phage HP1 and Slur09 that we retrieved from SRA. These phages were sequenced using illumina nextera kits which lead to the loss of the termini sequence. As expected no termini could be detected when using the data generated for these two phages.

All the PhageTerm results are detailed in Table 2 and reports for each phage are available in the sourceforge deposit (https://sourceforge.net/projects/phageterm).

**Table 2.** Summary of PhageTerm results

| Phage Name | Ends | Left | Right | Class | Type |
| --- | --- | --- | --- | --- | --- |
| **Phages references** |  |  |  |  |  |
| Lambda | Non Red. | Fixed | Fixed | COS (5') | Lambda |
| HK97 | Non Red. | Fixed | Fixed | COS (3') | HK97 |
| T7 | Red. | Fixed | Fixed | DTR (short) | T7 |
| P1 | Red. | Preferred | Distributed | Headful (pac) | P1 |
| T4 | Red. | Random | Random | Headful | - |
| Mu | Non Red. | Random | Random | Mu-like | Mu |
| **Phages of *C. difficile*** |  |  |  |  |  |
| phiCD481-1 | Non Red. | Fixed | Fixed | COS (3') | HK97 |
| phiCD506 | Non Red. | Fixed | Fixed | COS (3') | HK97 |
| phiMMP01 | Red. | Preferred | Distributed | Headful (pac) | P1 |
| phiMMP03 | Red. | Preferred | Distributed | Headful (pac) | P1 |
| phiCD211 | Red. | Fixed | Fixed | DTR (short) | T7 |
| phiCD146 | Red. | Preferred | Distributed | Headful (pac) | P1 |
| phiCD505 | Red. | Multiple | Multiple | - | - |
| phiCD111 | Red. | Multiple | Multiple | - | - |
| phiCD24-1 | Red. | Random | Random | - | - |
| **Phages from SRA** |  |  |  |  |  |
| T3 | Red. | Fixed | Fixed | DTR (short) | T7 |
| T7 | Red. | Preferred | Preferred | DTR (short) | - |
| Efm1 | Non Red. | Fixed | Fixed | COS (3') | HK97 |
| PBES-2 | Red. | Fixed | Preferred | DTR (short) | - |
| HP1 (nextera) | Red. | Multiple | Multiple | - | - |
| Slur09 (nextera) | Red. | Multiple | Multiple | - | - |

## DISCUSSION

PhageTerm provides an easy interface for biologists to decipher phage termini and mode of packaging using NGS data. Our strategy relies on the analyses of the number of reads starting at each position along the genome divided by the sequence coverage. This value that we term X has several informative properties that allows for the easy classification of phage types. This value is expected to be 0.5<X<1 for cos and DTR phages, and ~1/(C+1) for pac phages where C is the number of phage copies per concatemer. Depending on the number of significant peaks that are detected, their size and orientation, it is possible to classify phages according to 6 types: cos 3’, cos 5’, DTR short, DTR long, headful with no pac detected, headful with a pac site. An additional analysis of hybrid fragment carrying both phage and host DNA allows identifying phages that amplify through replicative transposition (Mu-like phages). When no termini are detected and the phage cannot be classified as Mu-like, PhageTerm currently does not make any prediction of the packaging mechanism. In such cases, if the user is confident that the phage genome provided is complete, it can likely be classified as a T4-like phage. However, we cannot exclude other possible mechanisms such as a phi29-like packaging strategy^15^ which involves proteins covalentely bound to the DNA termini. If proteins are not removed before adapter ligation during the sequencing library preparation, then we do not expect to obtain a signal with our strategy.

To compare our strategy with previous work, we also implemented in PhageTerm the approach described by Li *et al.*^17^. In this work, the authors analyze the number of reads starting at each position along the genome and classify the phages according to the number and orientation of peaks, their size and the ratio of the first peak to the second peak. In most cases both methods give identical results, although PhageTerm performs a more finegrained classification. A few cases highlight the benefits of using our method instead of the crude number of reads starting at the termini. Clostridium phage PhiCD146 has a duplicated region that was collapsed during the *de novo* assembly. This can be seen as a sudden increase of coverage in this region. Li’s method assigns a wrong terminus in this highcoverage region, while PhageTerm has no problem. In another example, Clostridium phage PhiCD481-1 is correctly identified as a 3’ cos phage by PhageTerm while Li’s method mistakenly classifies it as a pac phage missing a peak in the reverse orientation, where the coverage is unexpectedly low.

In previous work other strategies have been employed to identify packaging mechanism. In a recent publication by Rashid, J. *et al*.^30^ analyzing termini and packaging of *C. difficile* phages, researchers inferred packaging mechanism by looking at homologies between the terminase of newly sequenced phages and that of phages with known packaging mechanisms. This strategy only provides very partial and uncertain information about packaging mode and no information on termini type and sequences. Recently, another study made available a procedure to analyze whole genomes, physical ends and packaging strategies of phages using a pre-existing software^31^. This procedure can be very time consuming and contains numerous steps requiring bioinformatic skills, including the installation of the Phamerator program^32^ and the management of the related SQL databases. The heavy and unwieldy procedures available to phage scientists motivated our efforts to develop PhageTerm, an easy-to-use program that consolidates in a single straightforward analysis numerous highly valuable pieces of information about phage termini and packaging mode.

## CONCLUSIONS

PhageTerm provides a rapid and reliable analysis of NGS data to determine phage termini and packaging mode, provided that data is generated following some simple rules: (i) random fragmentation should always be used when preparing sequencing libraries. (ii) Paired-end sequencing should be used to obtain a more complete characterization of the termini.

The broad availability of phage termini information will in the future enable to shed light on the diversity of packaging mechanisms in nature. The PhageTerm software will also attempt to assign termini type even when these do not correspond to known phage packaging mechanism. As such the software might help in the identification of novel packaging strategies. We encourage the phage community to contact us in the case of such discovery and we will be happy to update the software to better take into account any novel class. Finally, similar approaches should be possible to determine the termini of Archaea and Eukaryotic viruses.

## AVAILABILITY AND REQUIREMENTS

Project name: PhageTerm

Project home page: https://sourceforge.net/projects/phageterm

Operating system(s): Platform independent

Programming language: Python

Other requirements: matplotlib 1.3.1, numpy 1.9.2, pandas 0.19.1, sklearn 0.18.1, scipy 0.18.1, statsmodels 0.6.1, reportlab 3.3.0.

License: General Public License version 3.0 (GPL3.0)

Any restrictions to use by non-academics: No

## AKNOWLEDGMENTS

The authors would like to thank Laurence Van Melderen and Ariane Toussaint for providing strain RH8816, Erica Lieberman from Eligo Bioscience for providing the sequencing data for phage HK97 and Laurent Debarbieux for providing phage T4 as well as useful discussions. We also acknowledge the Plate-forme Genomics and the center of informatics for Biology, especially Christiane Bouchier, Laurence Ma, Eric Deveau, Fabien Mareuil and Olivia Doppelt-Azeroual for their help and support with sequencing and Galaxy integration. This work was supported by a discovery grant from the Natural Sciences and Engineering Research Council of Canada (NSERC #341450-2010), the Agence Nationale de la Recherche (“CloSTARn”, ANR-13-JSV3-0005-01) and the french Government's Investissement d'Avenir program; Laboratoire d'Excellence ‘Integrative Biology of Emerging Infectious Diseases’ [ANR-10-LABX-62-IBEID].

## AUTHOR CONTRIBUTIONS

JG, MM and DB wrote the PhageTerm code. FD performed the sequencing preparation. MM, LCF and DB designed the study. JG, MM and DB wrote the manuscript. All authors have contributed to it and have read and accepted the final version.

